# Predicting bioactivity of antibiotic metabolites by molecular docking and dynamics

**DOI:** 10.1101/2022.09.06.506739

**Authors:** Hokin Chio, Ellen E Guest, Jon L Hobman, Tania Dottorini, Jonathan D Hirst, Dov J Stekel

## Abstract

Antibiotics enter the environment through waste streams, where they can exert selective pressure for antimicrobial resistance in bacteria. However, many antibiotics are excreted as partly metabolized forms, or can be subject to partial breakdown in wastewater treatment, soil, or through natural processes in the environment. If a metabolite is bioactive, even at sub-lethal levels, and also stable in the environment, then it could provide selection pressure for resistance. (5S)-penicilloic acid of piperacillin has previously been found complexed to the binding pocket of penicillin binding protein 3 (PBP3) of *Pseudomonas aeruginosa*. Here, we predicted the affinities of all potentially relevant antibiotic metabolites of ten different penicillins to that target protein, using molecular docking and molecular dynamics simulations. Docking predicts that, in addition to penicilloic acid, pseudopenicillin derivatives of these penicillins, as well as 6-aminopenicillanic acid (6APA), could also bind to this target. Molecular dynamics simulations further confirmed that (5*R*)-pseudopenicillin and 6APA bind the target protein, in addition to (5S)-penicilloic acid. Thus, it is possible that these metabolites are bioactive, and, if stable in the environment, could be contaminants selective for antibiotic resistance. This could have considerable significance for environmental surveillance for antibiotics as a means to reduce antimicrobial resistance, because targeted mass spectrometry could be required for relevant metabolites as well as the native antibiotics.

## 1. Introduction

Antibiotics are widely used in medicine and agriculture. Moreover, the most recent report from WHO showed that the global antibiotic consumption is increasing, from 21.1 billion defined daily doses in 2000 to 34.8 billion defined daily doses in 2015 (1). Orally administered antibiotics, whether in human or veterinary medicine, may be absorbed in the gut; and so may be partly or wholly metabolized prior to excretion, while unabsorbed antibiotics will exit via faeces. Intravenous antibiotics may be subject to similar metabolic fates in serum, faeces and urine (2). Most administered antibiotics are not digested: 50-90% of antibiotic intake is excreted and 30% of antibiotics are excreted as metabolites. However, fewer than a third of antibiotic metabolites have yet been tested for bioactivity (3–5). Antibiotics can also be found in the excreta in farm animals, but metabolite formation within farm animal excreta is largely unknown (6, 7).

Therefore antibiotics from medical and veterinary use, and their metabolites, enter the environment and appear as contaminants in wastewater, soil, surface and ground water, sewage, and wastewater treatment plants (8, 9). Antibiotic concentration can drive selection for antibiotic resistance. Importantly, even sub-lethal concentrations of antibiotics can also drive selection (10); the consequence is that antibiotic metabolites, that might be insufficiently potent to have clinical value, might still be able to drive selection for resistance, and so be environmentally important, especially if present in stable forms. While there are studies about how some beta-lactam antibiotics affect specific organisms under controlled photolytic conditions (11), there is limited knowledge of the impact of metabolites on bacteria that would normally be affected by the cognate antibiotic, nor on their environmental stability or bioactivity. Notably, however, one crystallographic study has found (5*S*)-penicilloic acid complexed to the binding pocket of penicillin binding protein 3 (PBP3) of *Pseudomonas aeruginosa* (12). This raises the important question as to whether a much wider range of antibiotic metabolites may also have this ability. Although wastewater treatment can reduce concentrations of antibiotics, it cannot completely eliminate antibiotics or their metabolites (11, 13, 14). Moreover, antibiotic and metabolite contamination may be greater in many lower and middle income countries, where wastewater treatment is limited.

Chemical detection of antibiotics is an important strand of effective surveillance against antimicrobial resistance. Antibiotics can be detected from the environment through high-performance liquid chromatography (HPLC), mass spectrometry (MALDI-TOF MS and LC-MS) and colorimetric sensor arrays (13, 14). However, these assays require knowledge of the chemical structure of the antibiotic to be detected and standardization (15): antibiotic metabolites will not be identified unless specifically tested for. If metabolites are important, then this is a possible surveillance omission.

In this study, we predict possible bioactivity of antibiotic metabolites using computational methods - molecular docking and molecular dynamics (MD) simulations - to predict binding of antibiotic metabolites to their target binding sites. We exemplify this method using the binding of penicillins to PBP3 of *Pseudomonas aeruginosa*(12), recognizing that the approach should be valid for other classes of antibiotics, to other target molecules, and in other organisms. Penicillins are chosen for three reasons. First, penicillins are important because they are the most commonly used antibiotic for human medicine (52% of Defined Daily Doses of total antibiotic used), as well as widely used in agriculture. Second, the chemical structure of penicillin metabolites are well known, because the degradation pathways of beta-lactams are well characterized (16). Third, the availability of crystal structures of both piperacillin and (5*S*)-penicilloic acid to to PBP3 (12), with resolution 2.31 Å, provides both an effective starting point for computational investigation of other metabolites within the penicillin family, as well as clear evidence that molecular interactions between metabolites and antibiotic targets are possible.

*Pseudomonas aeruginosa* is chosen because it is a Gram-negative bacterium on which beta-lactam antibiotics can have an impact. It causes disease in plants, animals and humans; most importantly, it causes serious infections in immunocompromised cancer patients, cystic fibrosis patients, and patients suffering from severe burns (17). Moreover, *P. aeruginosa* can be found widely in nature, in soil and water (18). There are three classes of *P. aeruginosa* PBPs, class A, B, and C (19). We have chosen PBP3 because PBP3 is a class B enzyme that performs transpeptidation and is inhibited by penicillin (20), while PBP4 is a class C enzyme, which only act as carboxypeptidases or endopeptidases. The key residues that stabilize the antibiotic-PBP interaction are known: Ser294, Ser349, Ser485, Thr487 and Tyr503 (12, 21). In particular, Ser294 is the main residue for the antibiotic-PBP interaction as the residue that forms the covalent bond with the beta-lactam ring (21).

Antibiotics in the penicillin family contain an unstable, highly strained and reactive beta-lactam amide bond (22). While the beta-lactam bond is essential for the clinical efficacy of penicillins, it is not known whether penicillin metabolites are sufficiently bioactive to have selective impact. The degradation of penicillin takes place in a wide range of conditions, both alkaline or acidic (Fig. 1) (23, 24), in the presence or absence of the enzyme ß-lactamase, or under the action of weak electrophiles including water and metal ions. Penicillin undergoes further isomerization to penicillenic acid. The beta-lactam ring and its amide bond break open in the presence of acid giving an array of products, including penilloic acid, penicillamine and penilloaldehyde, through intermediates, namely penillic acid, penicilloic acid and penicillenic acid. (16, 25). Metabolites can form in *R* and *S* stereoisomer forms, because the oxidizing agent can attack from the upper plane or the lower plane of the penicillin. These stereoisomers may have different effects on the affinity of the metabolites towards the PBP (26).

**Figure 1.**
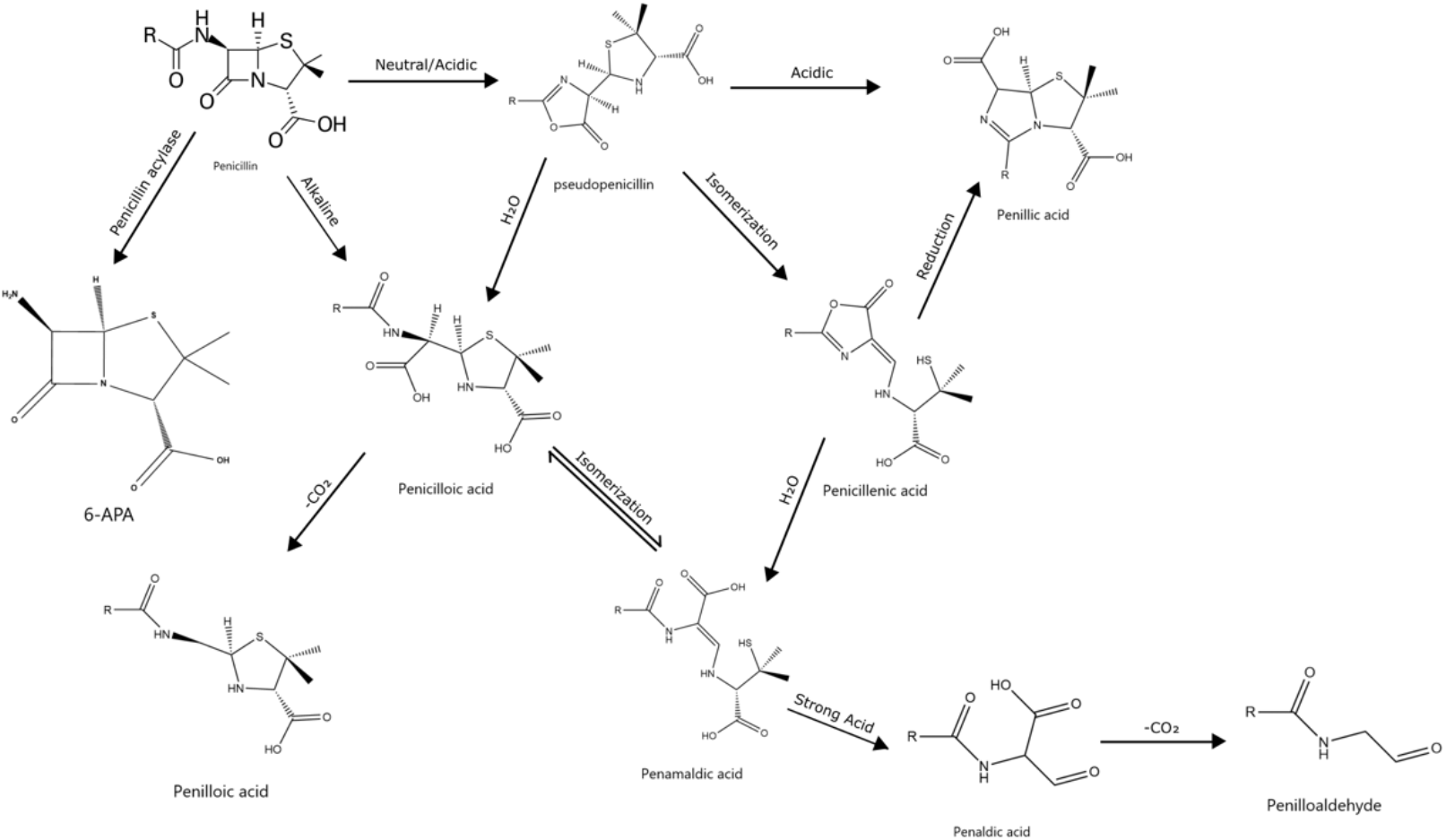
Pathways of the degradation of beta-lactam (penicillin family). The penicillin will degrade into 6-APA, pseudopenicillin, penicilloic acid, then further degrade to penilloic acid, penamaldic acid, penicillanic acid, penillic acid. The penamaldic acid further degrades to penaldic acid then penilloaldhyde under strong acid.

## 2. Methods

### 2.1 Docking

The reference protein structures used for docking were taken from X-ray structures deposited in the Protein Data Bank (www.rcsb.org). We used 4KQO, the crystal structure of PBP3 from *P. aeruginosa* in complex with piperacillin (12). The docking software, GOLD 5.7.1 (27), was used. All water molecules within the crystallized structure were removed and hydrogen atoms were added when missing from the PDB structure. For each protein target, the active site was defined as the collection of the amino acids enclosed within 8 Å radius sphere which calculated by eBoxSize (28) and suggests optimized box size x=y=z=15 Å, centered on the bound antibiotic ligand and flexible docking for 10° of movement freedom of the key residue (Ser294, Ser349, Ser485, Thr487 and Tyr503). The docking used the automatic genetic algorithm setting with 100% search efficiency. For the ligand flexibility, the internal hydrogen bonds were detected. The ring conformations, planar amide bond and protonated carboxylic acids were allowed to flip. The torsion angle distributions and postprocess rotatable bonds used the default parameter file. The fitness level was calculated by Piecewise Linear Potential method.

The molecular docking process was verified by redocking the ligand back into the crystal structure PBP3 complex with ligand removed. Nine further antibiotics (amoxicillin, ampicillin, cloxacillin, dicloxacillin, flucloxacillin, penicillin G, penicillin V, ticarcillin, oxacillin) were selected to create a reference result towards the docking of the 13 metabolites: (5*R*)- and (5*S*)-penicilloic acid, (5*R*)- and (5*S*)-penillic acid, (5*R*)- and (5*S*)-penilloic acid, (5*R*)- and (5*S*)-pseudopenicillin, penamaldic acid, penicillenic acid, penaldic acid, penilloaldehyde and 6-APA. The first step was to dock the antibiotic itself in order to produce a standard binding pose for that antibiotic. The second step was to dock each metabolite for that antibiotic which is compared with its standard binding pose. 13 metabolites of the antibiotics were screened by docking the molecules into the same binding pocket of the PBP. Ligand interactions were depicted using MOE 2015, while the 3D structures of the ligand with the binding site were visualized using PyMol 2.3.3.

The fitness was calculated as the sum of the steric complementarity between protein and ligand, the internal score of the ligand consists of the heavy-atom clash potential, the torsional potential, covalent docking and flexible sidechains (27). The RMSD of the docked metabolite was calculated by comparing the positions of the carbon atoms of the beta-lactam ring of the docked metabolite with their positions in the docked cognate antibiotic.

### 2.2 Molecular dynamics simulations

The quick MD simulator module in CHARMM-GUI (29) was used to add hydrogen atoms to the crystal structure, solvate the system with the TIP3P model (30), and apply periodic boundary conditions under the CHARMM force field (31). As one atomic angle parameter for the target protein (CG2O3-CG3C51-NG3C51) was not present in the force field settings, initial estimates for that angle were obtained using the CGenFF atom typing program (32, 33). The system was solvated in a truncated octahedral periodic boundary cell with edge distances 10 Å from the protein surface.

Energy minimization for each system was performed in NAMD 1.12 (34) using the standard conjugate gradient algorithm for 10,000 steps. All heating, equilibration, and production dynamics were performed using NAMD with a time step of 2fs, the CHARMM36 force field (35), and periodic boundary conditions. The parameters for the bile salts were taken from the CHARMM general force field (CGenFF) of drug-like small molecules.

The system was heated from 0 to 298 K in increments of 5K by temperature reassignment, where the velocities of all the atoms in the system are reassigned so that the entire system is set to the target temperature. The velocities were reassigned every 500 time-steps for 50,000 time-steps in the NVT ensemble. The systems were equilibrated for another 5ns in the NPT ensemble, with Langevin dynamics the pressure set to 1 atm. Production dynamics were run in the NPT ensemble for 30ns for each of the metabolites and the antibiotics. The equilibration was monitored by root mean square deviation (RMSD) trajectory analysis.

The analysis of the MD simulation is performed in VMD 1.94a51(36). The RMSD trajectory was calculated as the average RMSD of the protein including the ligand in each frame. The RMSD calculation was based on the backbone atoms for residues 250 to 477. Root-mean-square-fluctuation (RMSF) was calculated to quantify flexibility of the individual residues over the simulation. The interaction frequency between the ligand and each target residue was the proportion of 1000 frames in which the distance between the ligand that that residue was less than 3Å. The interaction frequency is also calculated for the systems with piperacillin, (5*R*)- and (5*S*)-penicilloic acid, *(5R)-* and (5*S*)- pseudopenicillin, penamaldic acid, 6APA and decoy molecules.

## 3. Results

### 3.1 The docking pose of piperacillin closely matches the crystal structure

The first step was to verify the integrity of the method by docking an antibiotic ligand back into its cognate binding site from a known crystal structure. We used crystallized piperacillin into 4KQO, PBP3 of *P. aeruginosa*. This produced a benchmark against which the binding of other antibiotics or metabolites to the PBP binding site could be compared. The docking results (Figure 2a) showed that the RMSD of the redocking of piperacillin is 0.62 Å with fitness 149. This was confirmed by the proximity of the redocked piperacillin to its position in the crystal structure (Figure 2b). This gave confidence that the result was a suitable benchmark for docking the antibiotics and metabolites.

**Figure 2.**
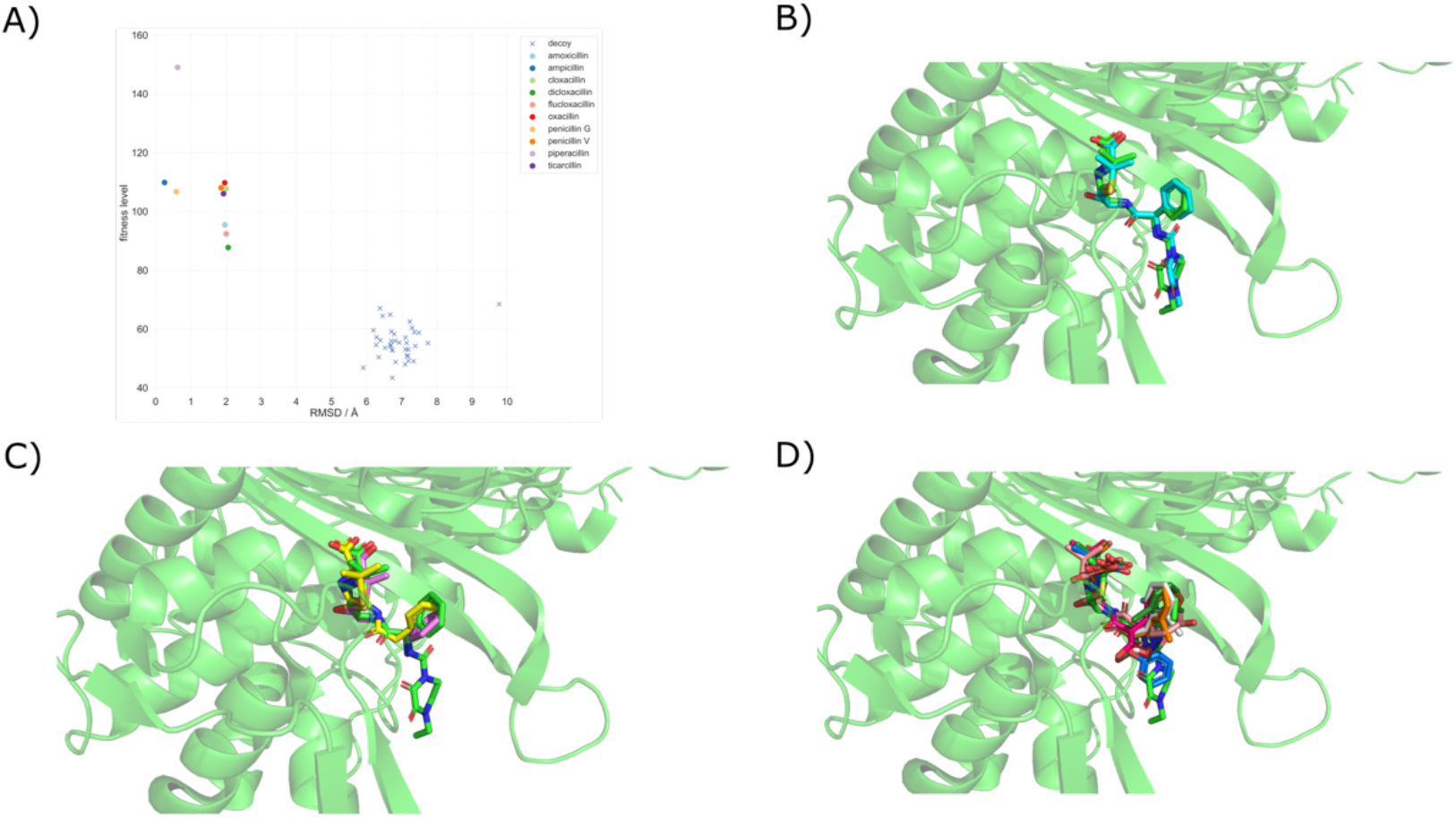
A) Docking results showing the fitness levels and RMSDs of antibiotics and decoys. B-D) Binding poses from docking of b) piperacillin (X-ray structure in green) and docked piperacillin (light blue), c) penicillin G (yellow), ampicillin (purple) which have lower RMSD, c) amoxicillin (pink), cloxacillin (white), dicloxacillin (dark green), flucloxacillin (orange), d) oxacillin (brown), penicillin V (blue), ticarcillin (dark red).

### 3.2 Docking of antibiotics

The same docking setup was used to dock the nine antibiotics (amoxicillin, ampicillin, cloxacillin, dicloxacillin, flucloxacillin, penicillin G, penicillin V, ticarcillin, oxacillin) into the PBP with the common beta-lactam ring as reference. Ampicillin and Penicillin G showed lower RMSD than piperacillin, with 0.25 Å and 0.58 Å, respectively (Figure 2c), but both had lower fitness scores of 109.9 and 106.7. The other penicillins had higher RMSDs (between 1.8 and 2.1 Å). Although these penicillins had a higher RMSDs, they were still close to the crystalized structure of piperacillin (Figure 2d).

### 3.3 Docking of decoy molecules to penicillin binding protein and metabolites to decoy protein

As controls, 50 decoy ligands were produced from the Directory of Useful Decoys (DUD) (37). The RMSD of the decoy ligands docked to PBP were between 6 Å to 8 Å (Figure 2a, Figure 3), considerably higher than any of the antibiotics. The fitness scores of the docking of decoy ligands with the PBPs was between 43 and 68. Taken together, this suggested that the decoy ligands did not bind the PBP and provided useful quantitative controls for metabolite docking below.

**Figure 3.**
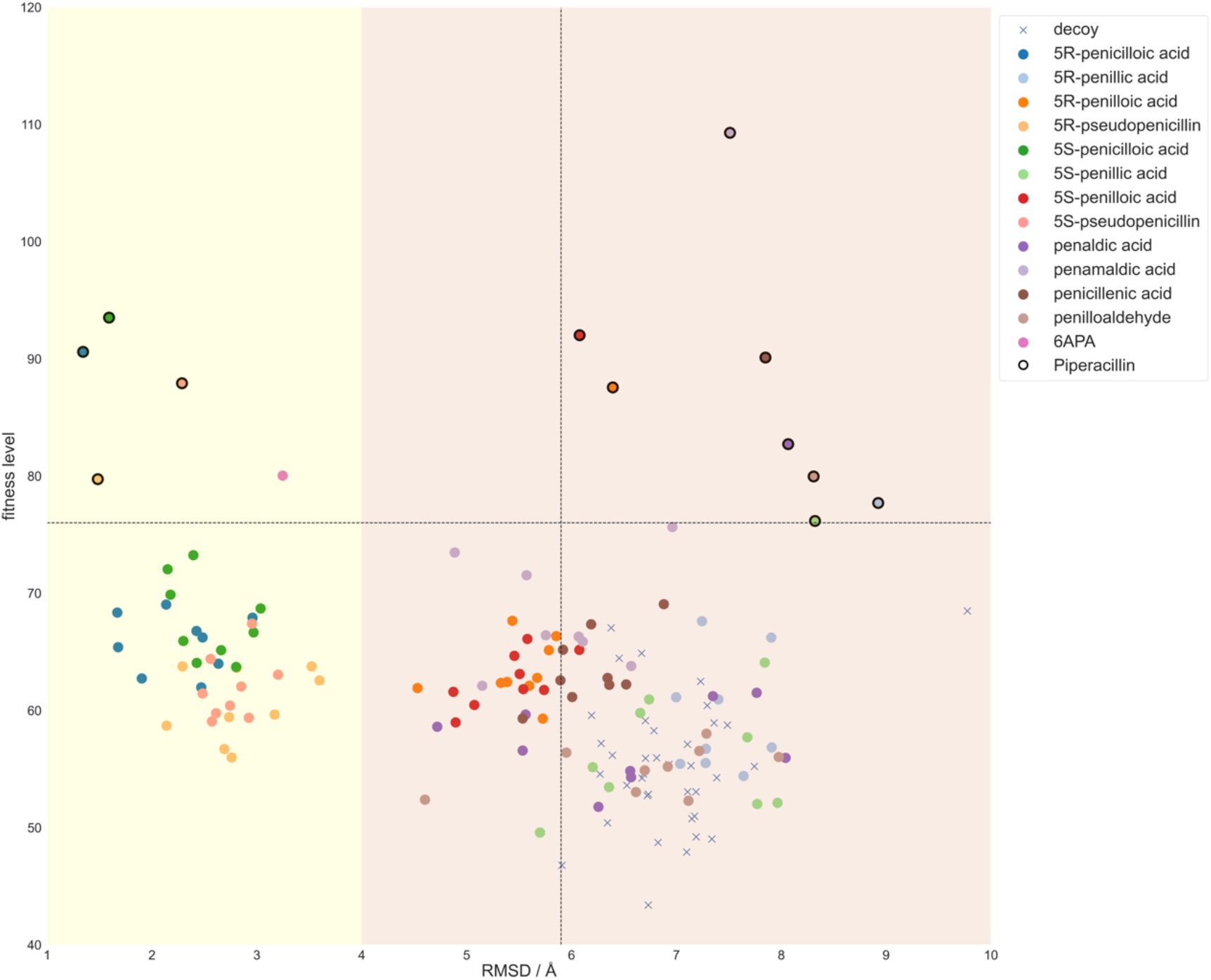
Docking results showing the fitness levels and RMSDs of antibiotic metabolites, together with decoys in order to indicate thresholds for likely docking. The two coloured regions separate the low RMSD (yellow) and high RMSD (red). The horizontal dotted line separates piperacillin metabolites from the metabolites of other antibiotics. As this PBP binds to piperacillin, the R group of piperacillin metabolite will be more familiar to bind to PBP and thus higher RMSD. The vertical dotted line indicates the decoy ligand with the lowest RMSD and acts as the threshold value to separate the difference between potential binding metabolites and the decoy.

We established thresholds for metabolite binding to PBP3 by docking metabolites of piperacillin to the decoy protein, as well as both metabolites and decoy ligands to two decoy proteins to which beta-lactam antibiotics would not be expected to bind: thrombin inhibitor (1BA8) and prohormone-processing carboxypeptidase (1AC5). The RMSD and fitness levels for decoy binding to decoy proteins (1BA8 and 1AC5) were between 5.32 Å and 9.76 Å and 32.1 and 65, respectively (Supplementary Figure 1). The distribution of RMSDs and fitnesses associated with decoy bindings suggested that good metabolite binding could be represented by RMSD less than 5.4 Å and fitness greater than 70. Specifically, antibiotic metabolites of piperacillin docked to the PBP had fitness scores between 77 to 110, while metabolites of docked to decoy proteins (1BA8 and 1AC5) had fitness scores below 74; similar fitness scores were observed for the decoy molecules docked to the decoy proteins (Supplementary figure 1).

### 3.4 Docking of metabolites with PBPs shows penicilloic acid, pseudopenicillin and 6APA are expected to bind to PBP

When docking the actual penicillin metabolites, (5*R*)- and (5*S*)-penicilloic acid, (5*R*)- and (5*S*)-pseudopenicillin and 6APA had a high fitness (79 to 93) and low RMSD (1.3 Å to 3.2 Å) (Figure 3). They were suggested to be more likely to bind to PBP3 compared with the other metabolites. These results are consistent with the X-ray crystal structure for PBP3 complexed with (5*S*)-penicilloic acid (12), providing confidence for the remaining predictions. (5*R*)- and (5*S*)-penilloic acid, penamaldic acid and penicillanic acid had a high fitness level (87.6 to 109) and high RMSD (6.1 Å to 7.8 Å) while the decoy molecules bound to PBP have similar RMSD (6 Å to 8 Å) and have a lower fitness (43 to 68) (Figure 3). While they might interact with PBPs, the pattern of interaction of the ligand might differ from the antibiotics so they were less likely to bind. (5*R*)- and (5*S*)-penillic acid, penaldic acid, penilloaldyhyde had low fitness level (76 to 82) and high RMSD (8.1 Å to 8.9 Å) and these molecules had a higher RMSD than the decoy molecules (Figure 3), so they could not bind to PBP or might easily detach from the protein after binding to it. Because penamaldic acid had the highest fitness and also a high RMSD, it had been chosen as a negative control for the MD simulations below. The metabolites docking to the PBP had fitness level between 77 to 110. When metabolites were docked to the decoy proteins, the fitness levels (37 to 74) were similar to the fitness level when the decoy molecules docked to the decoy proteins (Supplementary figure 1).

### 3.5 Predicted interactions of metabolites with key PBP3 residues

In order to verify the structural plausibility of the low RMSD predicting docking poses, we compared the 3D orientation of penicilloic acid and pseudopenicillin of piperacillin, as well as 6-APA, with that of the crystal structure for docked piperacillin. The antibiotic piperacillin interacts with the residue Ser294 with a covalent bond, and with residues Ser349, Ser485, Thr487 and Tyr503 with hydrogen bonds (Supplementary Figure 2). In comparison, (5*R*)-penicilloic acid interacted with Ser485 and Thr487, (5*S*)-penicilloic acid interacted with Ser294, Ser349 and Thr487. (5*R*)-pseudopenicillin interacted with Ser349, Ser485 and Thr48, (5*S*)-penicilloic acid interact with Ser294. (5*S*)-pseudopenicillin interacted with Phe533, penamaldic acid interacted with Thr487 and 6-APA interacted with Ser294 and Ser349 (Table 1); these predicted interactions are all with hydrogen bonds (Supplementary Figure 2). Thus each of the high fitness and low RMSD metabolites was predicted to interact with PBP3 with at least one residue in common with piperacillin.

**Table 1.**
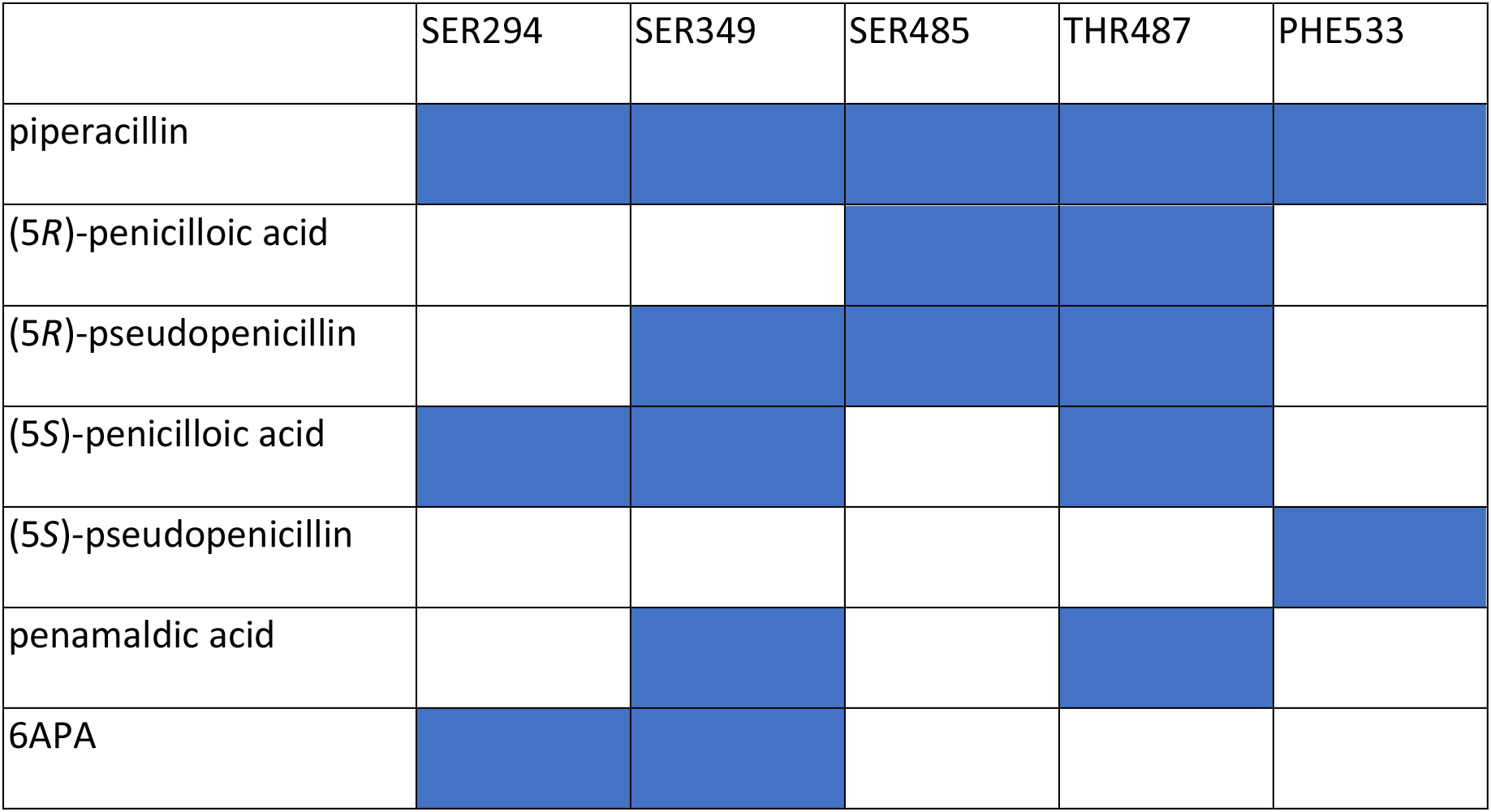
The filled boxes indicate interactions between residues of the PBP and antibiotics or metabolites based on the structural interaction between the ligand and the protein in supplementary figure 1.

### 3.6 MD simulations

In order to further assess potential for metabolites to bind into the PBP pocket, we ran MD simulations for the highest scoring metabolites (5*R*)- and (5*S*)- penicilloic acid, (5*R*)- and (5*S*)- pseudopenicillin and 6APA), with piperacillin as a positive control, and three negative controls: penamaldic acid, a decoy ligand with low fitness score, and a decoy ligand with high fitness score. The MD simulation outcomes were subjected to further analysis: root-mean-square fluctuation (RMSF) to assess whether structure of the system was in equilibrium; interaction frequencies between the metabolite and binding pocket atoms; and the position and structural orientation of the metabolites when they were sitting within the binding pocket.

#### 3.6.1 RMSF analysis of the MD simulations

During the MD simulations, the structure and the RMSD of the backbone atoms relative to the initial structure of each frame remained stable when the complex is in equilibrium. RMSD trajectory analysis was performed on the binding pocket (between residue 250 and residue 504), as the residues not associated with the binding pocket had very high RMSDs, which adversely bias the average RMSD when calculating RMSD for the whole frame (Supplementary figure 3). The RMSDs in the binding pocket varied between 0.5 Å to 2 Å, indicating that the systems are stable.

The root-mean-square fluctuation (RMSF; Fig. 4a) showed the deviation of the position of a particle or residue with respect to an initial position over all frames. The binding pocket (residues 250-504) was shown to be stable because it had a low RMSF ranging between 0.5 Å to 1.5 Å (Figure 4b). The N- and C-termini of the protein have high RMSF, suggesting that the structure of the N- and C-termini moved considerably across all frames; thus these termini might affect the value in RMSD analysis when determining whether the system was stable. MD simulations of piperacillin showed that the key binding residues (Ser294, Ser349, Ser485, THR487 and TYR503) were all close to their positions and orientations in the crystal structure (RMSD of 1 Å) (Supplementary figure 3), giving confidence in the MD simulations (Figure 4c).

**Figure 4.**
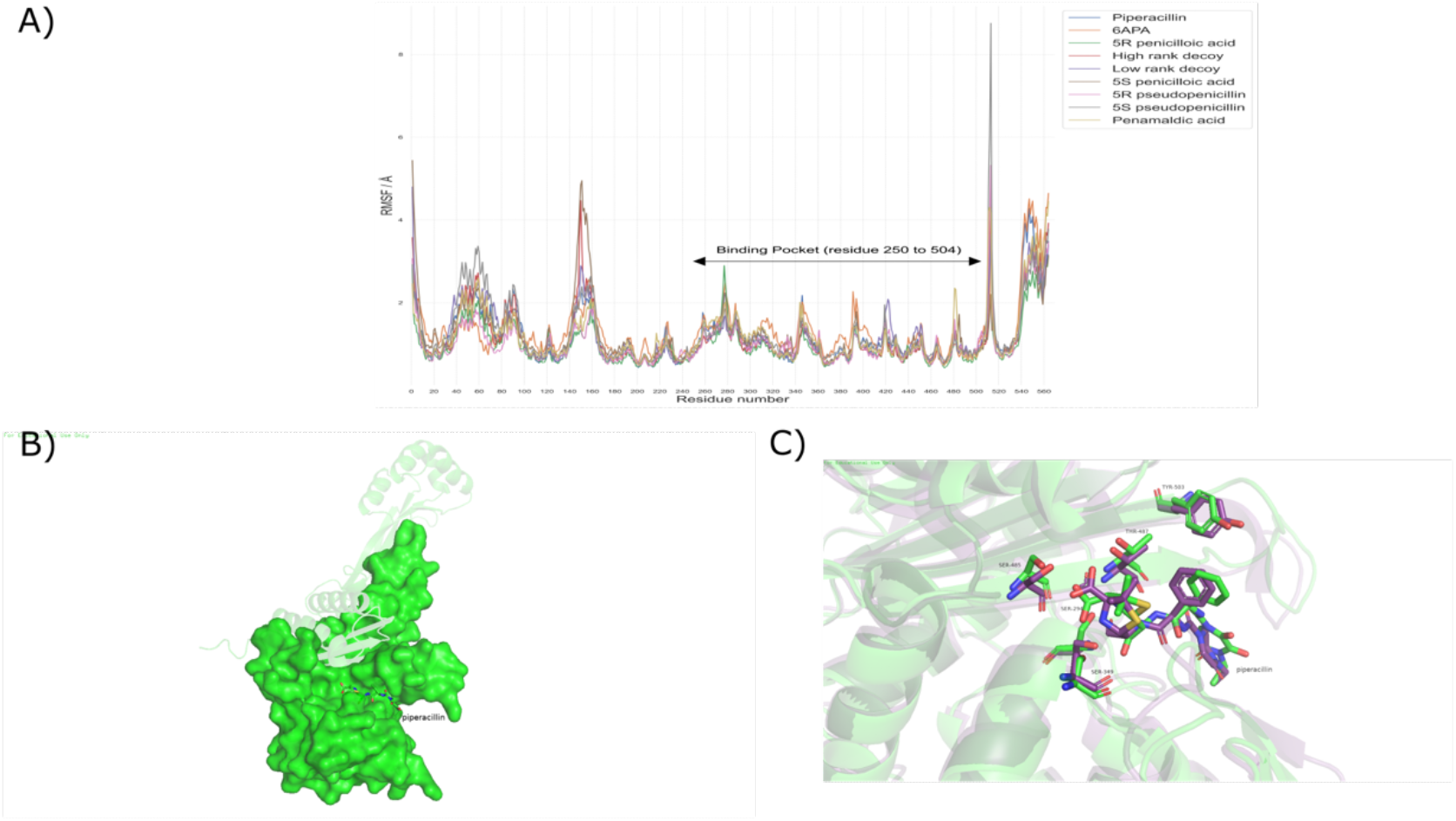
a) RMSF analysis of all MD simulations. It shows the RMSF of each residue among all frames. The binding pocket of the protein (residue 250 to 504) have a relatively fixed 3D structure in the binding pocket (RMSF below 2 Å), with the areas with highest RMSF lying outside of the binding pocket. b) the structure of binding pocket suggested by RMSF analysis indicated by the green surface, c) the orientation of the key residues and antibiotics piperacillin of MD stimulation (green) and the original pdb file (dark purple) showing good alignment between the MD simulations and crystal structure.

#### 3.6.2 Interaction frequency of the metabolites with the residues

Piperacillin interacted with the protein and was stabilized by the residues Ser294, Ser349, Ser485, Thr487 and Tyr503, with very high frequency. In order to assess binding of metabolites in more detail, the interactions between these residues and the ligands were assessed, by counting the frequency of distance below 3 Å across all frames (Table 2). The interactions of (5*R*)-pseudopenicillin and 6APA with key residues was close to that of piperacillin, with ligand-protein interactions at four of the five residues. (5*S*)-penicilloic acid and (5*S*)-pseudopenicillin were predicted to interact with two residues, while (5*R*)-penicilloic acid and penamaldic acid interacted at just one residue. The decoy ligands showed relatively weak interactions with the protein, with one residue and two residues respectively.

**Table 2.** Interaction frequencies between molecules and PBP residues as predicted by MD simulations and docking. Percentage is the frequency in which residue is within 3 Å from the ligand in MD simulations; interactions below 3 Å are predicted to be strong enough to stable the ligand. Colours represent synthesis of MD and docking results: red are interactions predicted by both simulation methods; yellow are interactions predicted from MD; blue are interactions predicted from docking; no colour signifies no predicted interactions. Overall, (5*R*)-pseudopenicillin, (5*S*)-penicilloic acid and 6APA are predicted to bind to the PBP3 binding pocket.

|  | SER294 |  | SER349 |  | SER485 |  | THR487 |  | PHE533 |
| --- | --- | --- | --- | --- | --- | --- | --- | --- | --- |
| piperacillin | 100.0% |  | 100.0% |  | 99.5% |  | 100.0% |  | 75.4% |
| (5R)- penicilloic acid | 35.6% |  | 11.7% |  |  |  | 89.4% |  | 49.0% |
| (5R)- pseudopenicillin | 99.8% |  | 2.0% |  | 99.7% |  | 99.3% |  | 89.8% |
| (5S)- penicilloic acid | 82.9% |  | 40.3% |  | 0.5% |  | 98.2% |  | 47.9% |
| (5S)- pseudopenicillin | 67.7% |  | 43.5% |  | 0.4% |  | 90.9% |  | 88.3% |
| penamaldic acid | 31.0% |  | 30.0% |  | 21.7% |  | 89.7% |  | 32.8% |
| 6APA | 100.0% |  | 100.0% |  | 99.6% |  | 100.0% |  | 73.4% |
| high rank decoy | 98.1% |  | 41.4% |  |  |  | 31.9% |  | 34.2% |
| low rank decoy | 99.6% |  | 99.7% |  | 17.6% |  | 37.1% |  | 13.2% |

#### 3.6.3 Position and structural orientation of the metabolites when they bound to PBP

(5*S*)-penicilloic acid, (5*R*)- and (5*S*)-pseudopenicillin, 6APA located on the center of the binding pocket (Figure 5a) and so had the potential to interact with the key residues. (5*R*)-penicilloic acid (3.1 Å below piperacillin) and penamaldic acid (on right of piperacillin with 6.9 Å) were located on the right side of the binding pocket, while the low rank decoy was in the left (with 4.4 Å); none of these molecules were predicted to enter the binding pocket (Figure 5b). The high rank decoy entered the binding pocket but the orientation of the ligand was different from piperacillin (3.5 Å difference between the center of decoy and pipercillin) (Figure 5b), so the molecule was not predicted to interact strongly with the protein as they had only one residue and two residue interactions (Table 2). (5*R*)-penicilloic acid and penamaldic acid lacked the interaction of the key residue Ser294 (Table 2), while the decoys interacted with Ser294 but with different location in the binding pocket relative to piperacillin (Figure 5b).

**Figure 5.**
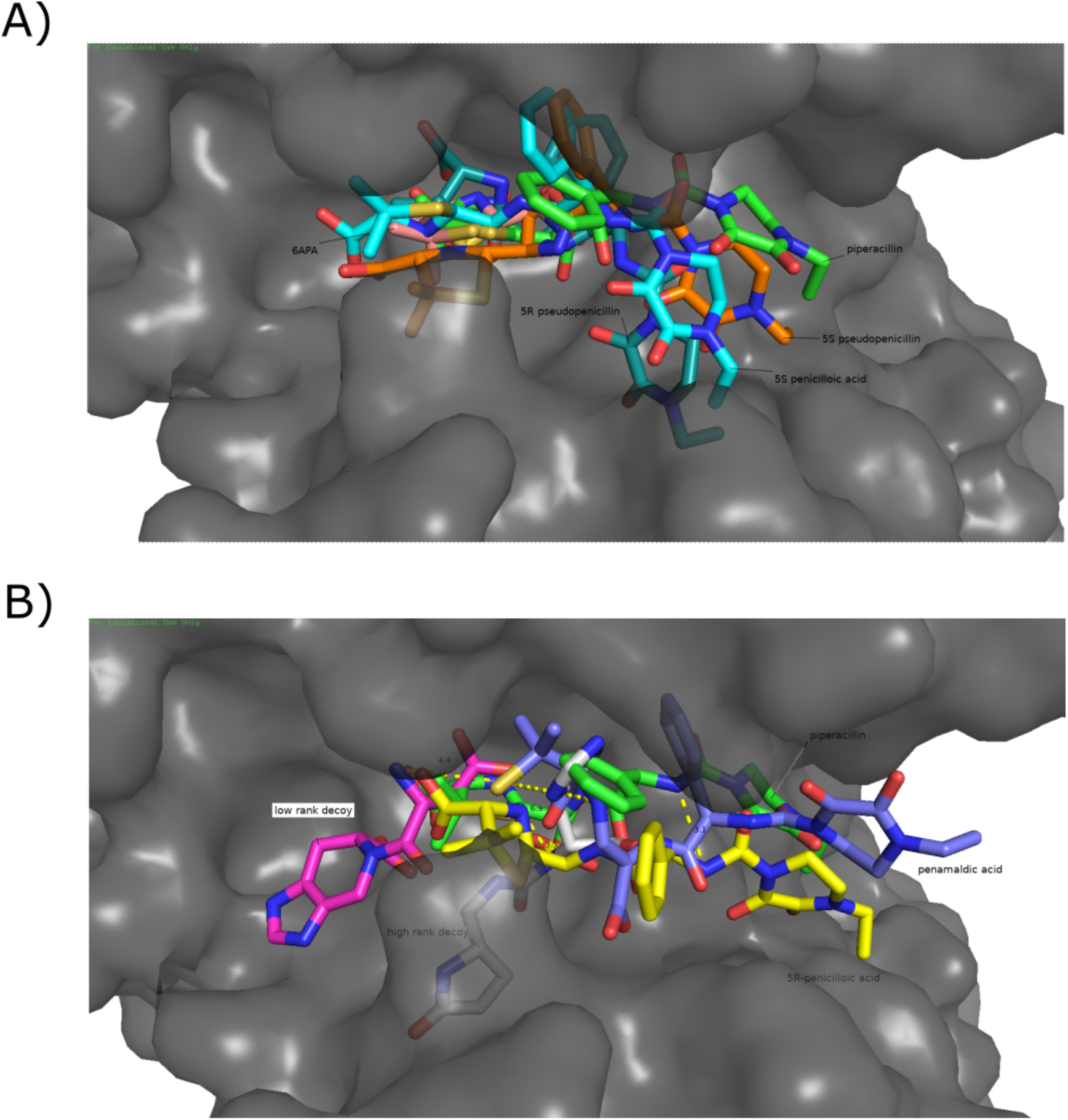
The orientation of the ligand within the protein. A) Piperacillin (green), (5*S*)-penicilloic acid (light blue), (5*R*)-pseudopenicillin (dark blue), (5*S*)-pseudopenicillin (orange) and 6APA (pink) sit in the binding pocket of PBP. B) (5*R*)-penicilloic acid (yellow) and penamaldic acid (blue). (5*R*)-penicilloic acid (3.1 Å below piperacillin) and penamaldic acid (on right of piperacillin with 6.9 Å) locate on the right side of the binding pocket and do not fully enter it. The low rank decoy (purple) is in the left (4.4 Å difference between the center of the decoy and piperacillin) and it is not in the binding pocket. The high rank decoy enters the binding pocket but it binds differently from the piperacillin (3.5 Å difference between the center of decoy and pipercillin).

We next compared the orientation and alpha carbon positions of the protein residues between metabolite and piperacillin binding (Figure 6 and Table 3). (5*R*)-psudopenicillin, (5*S*)-penicilloic acid and 6APA had a similar pattern of interaction of key residues and torsion angles as piperacillin, further confirming the prediction that they bound on the PBP. There was a large difference in RMSD of the alpha carbon of (5*R*)-(4.1 Å) and (5*S*)-(2.5 Å) pseudopenicillin on Ser349 (Table 3). The residue Ser349 moved outwards, and the binding pocket became larger. These might cause the interaction (2%) between the pseudopenicillin and the protein became weaker but also the movement of the alpha carbon of Ser349 was to adopt difference shape of pesudopenicillin and to allow the binding of psudopenicillin towards PBP. The torsion angles of residue among systems of (5*R*)-psudopenicillin, (5*S*)-penicilloic acid and 6APA showed they were similar to piperacillin (Figure 6).

**Figure 6.**
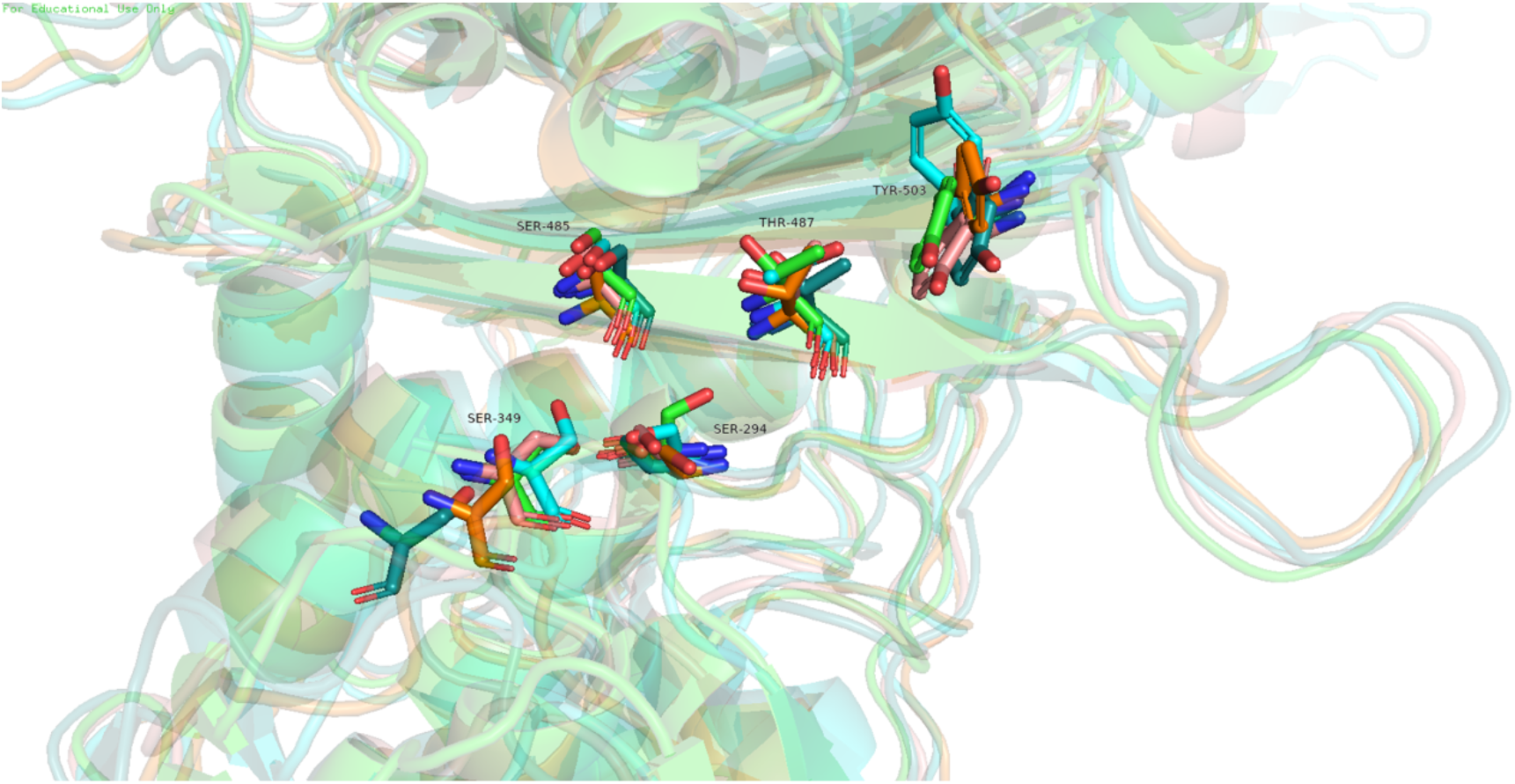
Structural orientations of the residues Ser294, Ser349, Ser485, Thr487 and Tyr503 in MD stimulations of piperacillin (green), (5*R*)-(dark blue) and (5*S*)- (orange) pseudopenicillin, (5*S*)-penicilloic acid (sky blue) and 6APA (pink) showing the movement of the alpha carbon and the torsion angle of the metabolites when comparing with the piperacillin (Table 2). The movement of the alpha carbon of the Ser349 of pseudopenicillin makes a larger binding pocket when compared with the binding pocket of piperacillin.

**Table 3.** Table of torsion angle of the residue and distance move of the alpha carbon of the metabolites in Å compared with piperacillin. The box filled with yellow shows there is a large change in the alpha carbon atom of the residue and this enlarged the binding pocket making it not likely to interact with the ligand.

|  | SER294 |  | SER349 |  | SER485 |  | THR487 |  | TYR503 |  |
| --- | --- | --- | --- | --- | --- | --- | --- | --- | --- | --- |
|  | Psi | Phi | Psi | Phi | Psi | Phi | Psi | Phi | Psi | Phi |
| Piperacillin | -52.2° | -27.6° | 30.3° | 66.5° | 159.4° | -93.5° | 108.7° | -125.7° | 160.7° | -132.6° |
| (5R)-<br>pseudopenicillin | -10.9° | -72.2° | 156.3° | 65° | 156° | -81.3° | 133.7° | -141.9° | 165.3° | -141° |
| (5S)-penicilloic acid | -12.1° | -63.6° | 17.1° | 60.5° | 166.8° | -99.7° | 143.1° | -144.5° | 154.9° | -159.9° |
| (5S)-<br>pseudopenicillin | -3.5° | -76.3° | 48.3° | 42.5° | 152.8° | -96.1° | 153.1° | -122.9° | 152.9° | -135.3° |
| 6APA | -1.4° | -70.2° | 26.2° | 82.9° | 160.9° | -69.6° | 119.6° | -155.9° | 156.6° | -129.8° |

|  | Distance of alpha carbon movement (Å) |  |  |  |  |
| --- | --- | --- | --- | --- | --- |
|  | SER294 | SER349 | SER485 | THR487 | PHE533 |
| (5R)-<br>pseudopenicillin | 0.8 | 4.1 | 0.9 | 1.5 | 0.8 |
| (5S)-penicilloic acid | 0.4 | 0.8 | 0.5 | 0.9 | 0.4 |
| (5S)-<br>pseudopenicillin | 0.8 | 2.5 | 1.2 | 0.9 | 0.5 |
| 6APA | 0.3 | 0.4 | 0.4 | 1.0 | 0.9 |

## 4. Discussion

In this study, docking predictions suggested that the metabolites (5*R*)- and (5*S*)-pseudopenicillin, (5*R*)- and (5*S*)-penicilloic acid and 6APA could bind PBPs. Through MD simulations, we see consistent predictions of possible binding of (5*R*)-pseudopenicillin, (5*S*)-penicilloic acid and 6APA into the binding pocket of PBP3. (5S)-penicilloic acid has already been found complexed to PBP3 in a stable crystal structure (12), lending confidence to our predictions for (5R)-pseudopenicllin and 6APA. In the MD simulations of (5*R*)-pseudopenicillin, four of the five main binding residues are in similar positions as compared with the MD simulation of penicillin (Table 3), with very high probabilities of interaction (Table 2). The most important residue, Ser294, interacting 99.8%, with its carbon atom moving by only 0.8 Å between the simulations of the two ligands. The exception is Ser349, whose alpha carbon moves by 4.9 Å. Although the position of Ser349 moved to adopt the shape of (5*R*)-pseudopenicillin, Ser349 did not interact with (5*R*)-pseudopenicillin (2%), so this may slightly weaken the interaction of the PBP with (5*R*)-pseudopenicillin. This suggests that molecules have the ability to bind to the PBP not only though the covalent bonding between serine and the oxygen of the nitrogen quadrilateral ring but also though the non-covalent interaction between the molecules and the penicillin binding protein (12). The data also show that the non-covalent binding (5*S*)-penicilloic acid and the antibiotic bind to the same binding site with similar orientation. Taken together, this gives confidence that these metabolites have the potential to bind to the target penicillin binding protein. If so, they may also and have biological effect, specifically providing some (non-clinical) level of antibiotic action, but possibly sufficient to provide selective pressure for resistance.

Penicillins are relatively unstable in the environment, e.g. piperacillin can remain stable with 5% tazobactam sodium for 2 days at 25°C (38), amoxicillin remains stable with clavulanic acid for 26 hours at 25°C (39) and penicillin G could be stable for 24 hours at 25°C (40). Although there is little evidence about the stability of penicillin metabolites in laboratory or environmental conditions, the penillic acid, penicilloic acid and penicillenic acid can be highly unstable metabolites with an unstable beta-lactam ring (25), so the evidence for the uncertain stability of the metabolite is required. Any environmentally stable antibiotic metabolites could be important contaminants, as low antibiotic concentrations may increase genetic variability of microbes and select for resistance (10, 41). This could have considerable significance for environmental surveillance for antibiotics, as stable metabolites that exert selective pressure for antibiotic resistance would also need to be measured.

PBPs are classified into high molecular mass and low molecular mass. High molecular mass PBPs are classified by the number of reactions that they can catalyze, class A and B (19). The bifunctional enzyme (class A) catalyzes both the glycosyltransfer and transpeptidation, while the monofunctional enzyme (class B) only perform transpeptidation (42). They are serine acyltransferases which catalyze the formation of cross-link peptidoglycan which is an essential macromolecule surrounding the bacteria. The low molecular mass PBPs (class C) generally function as carboxypeptidases or endopeptidases and typically be genetically deleted without having a significant effect on cell viability or growth (43). The PBPs are numbered in the order of decreasing molecular weight in a given organism, so there is no relationship between the same numbered PBPs of two unrelated organisms (e.g. PBP-2 of *E. coli* and PBP-2 *of P. Aeruginosa)*. The PBPs used in the study is PBP3 of *P*.

*Aeruginosa* which is class B PBP(19). This lays the ground for further detailed simulations of metabolite binding to other PBP classes (class A or class C), or in other organisms. Moreover, it would also be possible to use these docking and MD simulations to screen mutations of PBPs for antibiotic binding, and so predict mutational resistance to penicillins or their metabolites.

While we focused on penicillins, these methods could also be applied to the same antibiotic class but a different family, such as cephalosporins, or to other antibiotic classes, such as tetracyclines, aminoglycosides. Cephalosporins also target PBP and have same binding mechanism with penicillin as they belong to the beta-lactam family, but have different degradation pathways from penicillin and so have different metabolite products (16, 44). Tetracyclines and aminoglycosides both target the 30S ribosome subunit, but they bind to different positions of the ribosome. Tetracycline binds on the A-site of the 30S ribosome preventing the binding of the aminoacyl-tRNA to the A-site (45) and further preventing the formation of the protein. Aminoglycosides bind to the start codon of the 30S ribosome and form a complex to prevent the start of the formation of protein (46). There is detailed information of the degradation pathway of tetracycline (47), but the degradation of the aminoglycoside is less well characterized (48), thus placing tetracycline as more amenable for similar analysis.

An important limitation of our results is that we only had access to one structure of antibiotic complexed to its target protein in a single organism (4KQO). While the predictions have been successful, their generality would be improved with access to further structures of different PBPs, different penicillins, or different organisms.

Moreover, molecular docking cannot predict the presence of unknown covalent-binding (27), which may affect the predictions of how the ligand binds to the target protein. MD simulations also have limitations (35), with results that can depend upon the parameter sets used (49), and the molecular mechanics force field chosen (32). The free binding energy was not calculated in this study but it could be calculated though MM/PBSA and MM/GBSA (50). The uptake of the metabolites of the organisms also need to be estimated; antibiotics with cytoplasmic or ribosomal targets need to enter the cell, including crossing the cell wall and membrane, in order to perform their functions (51). This step is untested in our analysis of metabolite docking. The metabolic state of microbes may influence the antibiotics susceptibility of the microbes and this may affect the amount and effect of uptake of antibiotics (52). Finally, some details of the degradation pathways of penicillin are not yet fully understood; these may produce other metabolites which have not been tested in this study (53).

## 5. Conclusion

In this study, we predicted that (5*R*)-pseudopenicillin and 6APA can bind to their cognate protein and may be bioactive, in addition to (5*S*)-penicilloic acid which has already been shown to be binding to PBP3 (12). If these molecules are stable and present in the environment, then they could act as selective agents for antimicrobial resistance. Empirical testing of these predictions is essential because there is currently is evidence about the stability of any of these metabolites under environmental or laboratory conditions. If this is found to be the case for any of these metabolites, then these metabolites should be included in environmental testing for antibiotics. Further study on metabolites of other antibiotic classes is also implicated.

## Author Contributions

**Hokin Chio**: Investigation, Methodology, Software, Writing – Original Draft. **Ellen E Guest**: Methodology, Resources. **Jon L Hobman**: Supervision, Writing – Reviewing & Editing. **Tania Dottorini**: Supervision, Writing – Reviewing & Editing. **Jonathan D Hirst**: Resources, Supervision, Writing – Reviewing & Editing. **Dov J Stekel**: Conceptualization, Supervision, Writing – Reviewing & Editing.

## Declaration

The authors declare that they have no conflicting interests.

## Acknowledgments

We thank Hong Chen of Zhejiang University for discussion that led to this project and David Scott for helpful feedback on early drafts. MD simulations were carried out using the University of Nottingham’s Augusta HPC service. JDH is supported by the Royal Academy of Engineering under the Chairs in Emerging Technologies Scheme, grant number CiET2021_17.

## Notes

### Competing Interest Statement

The authors have declared no competing interest.

