## Supplementary Figures for "Predicting bioactivity of antibiotic metabolites by molecular docking and dynamics"

### Supplementary Data

Figure S1

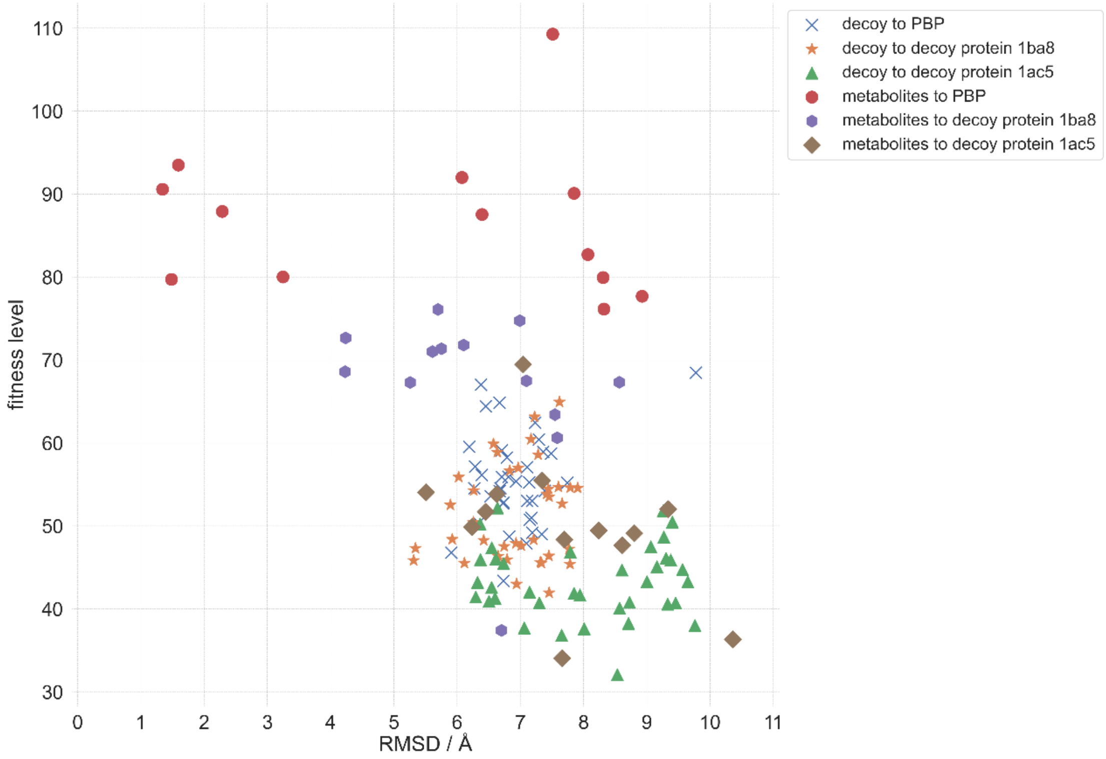


Figure S1. Docking results showing the fitness scores and RMSD of the metabolites and decoy ligands towards PBP and decoy proteins 1BA8 and 1AC5. The decoy ligands towards both PBP and decoy proteins have low fitness and high RMSD. The metabolites towards decoy proteins have slightly higher fitness scores than the decoy ligands and lower fitness scores than metabolites towards PBP and similar RMSD to decoys.

Figure S2

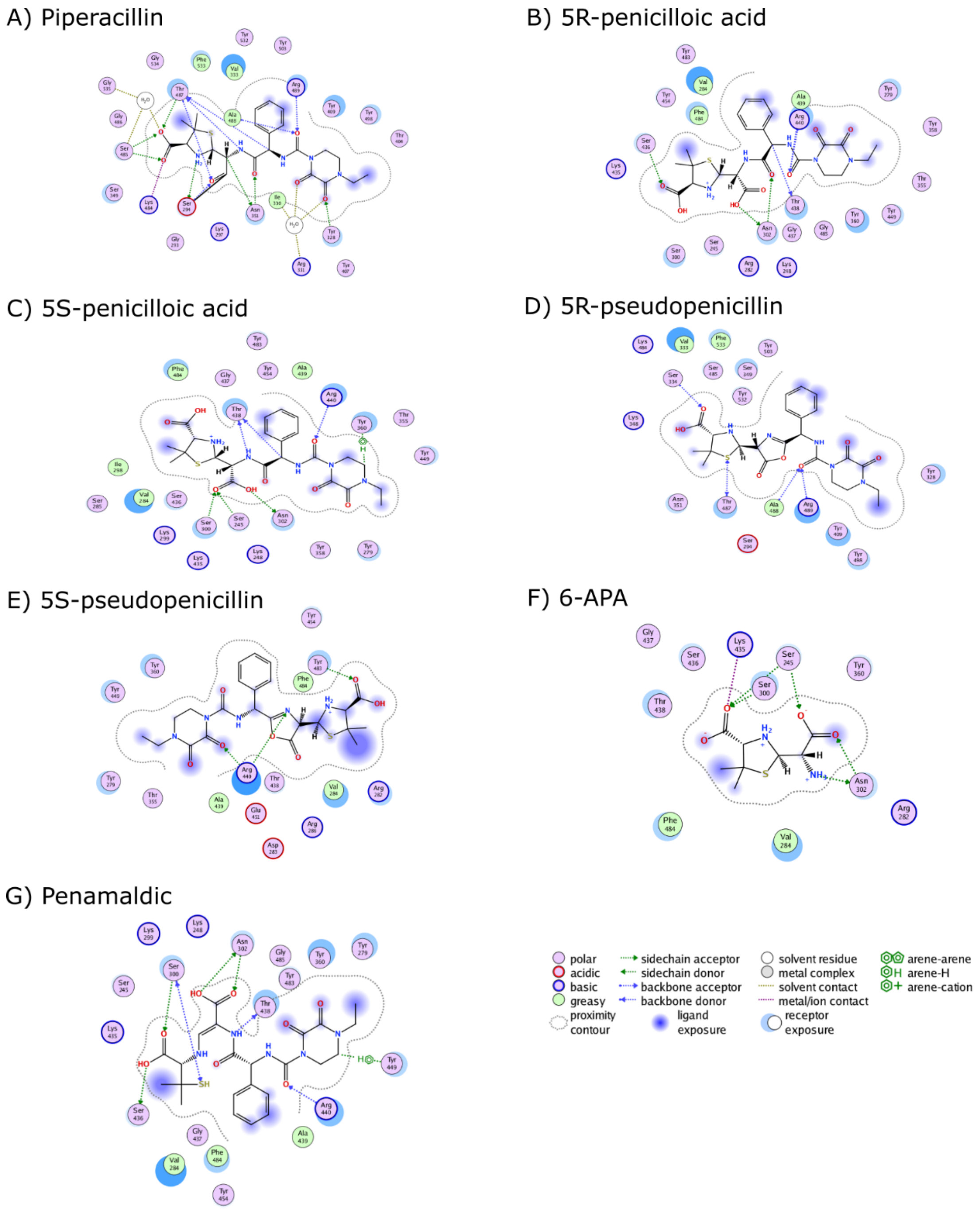


Figure S2. The interaction between the residues of the penicillin binding protein and A) piperacillin, B) (5*R*)-penicilloic acid C) (5*S*)-penicilloic acid, D) (5*R*)-pseudopenicillin, and E) (5*S*)-pseudopenicillin, F) 6APA and G) penamaldic acid of piperacillin with indication of the graph.

Figure S3

A)

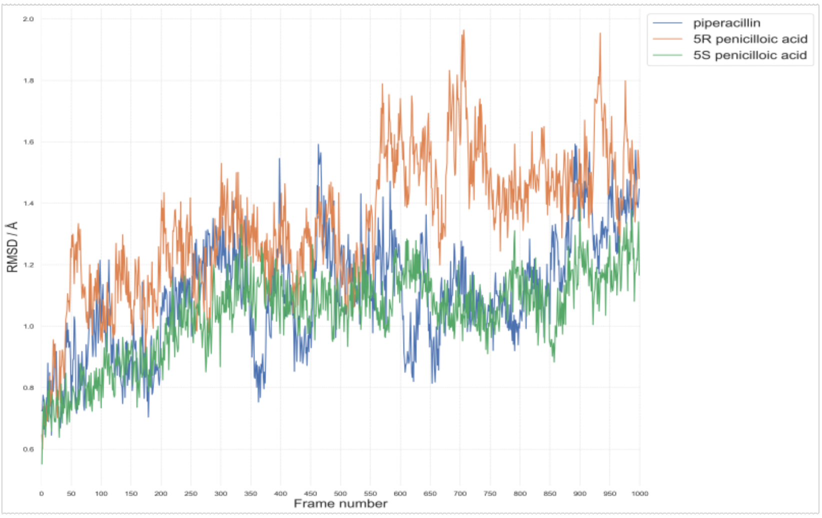


B)

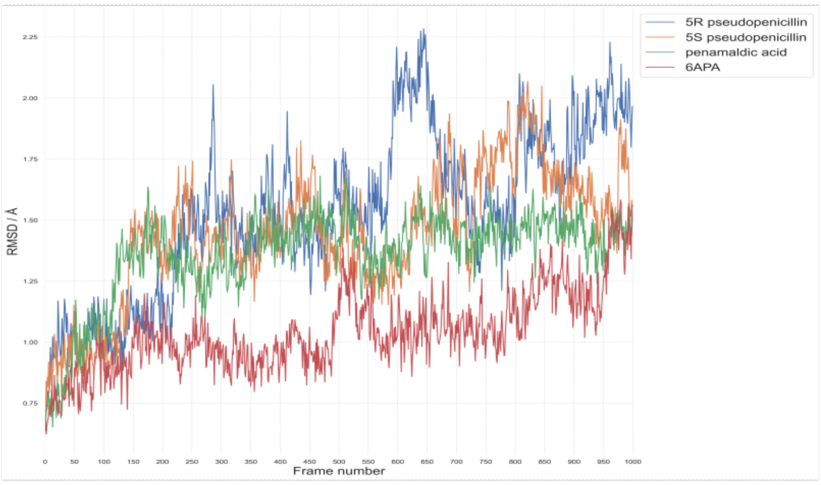


C)

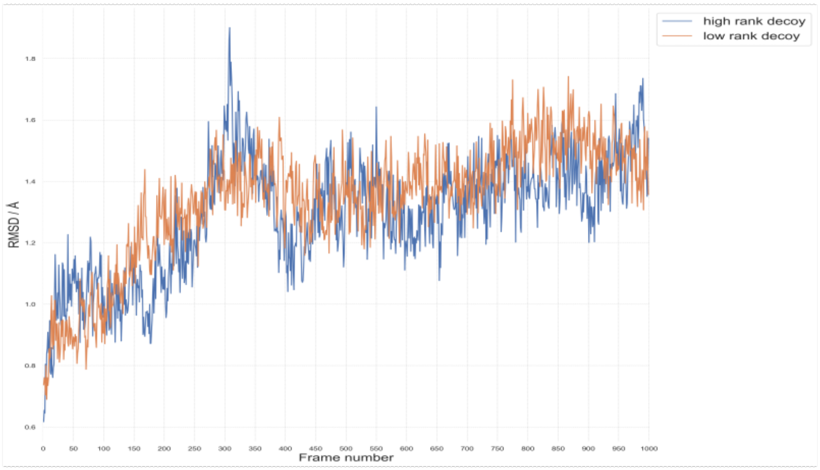


Figure S3a-d) RMSD of a) piperacillin, (5*R*)-penicilloic acid and (5*S*)-penicilloic acid, b) (5*R*)-pseudopenicillin, (5*R*)-pseudopenicillin, penamaldic acid and 6APA, c) decoys of the binding pocket (residue 250-477). The RMSD shows the system is relatively stable with their low RMSD.
